# Quaternary ice ages shaped protists phylogeography: the case of Arcellinida in the Iberian Peninsula

**DOI:** 10.1101/2025.08.11.669755

**Authors:** Rubén González-Miguéns, Emilio Cano, Enrique Lara

## Abstract

The Quaternary glaciations profoundly shaped the biogeography of plants and animals, yet their impact on microbial eukaryotes remains largely unexplored. We tested the “genetic legacy of the Quaternary” (GLQ) paradigm in terrestrial protists using Arcellinida testate amoebae diversity distribution across the Iberian Peninsula, a well-established glacial refugium. To do so, we compiled the most extensive Arcellinida metabarcoding dataset to date (ArKOI), including 615 samples from multiple continents and ecosystems. Our results showed that hotspots of intra-OTU genetic diversity align with known Iberian refugia, supporting the concept of “refugia within refugia”, and displaying clear ecoregion-specific climatic niches. Patterns of spatial clustering, niche breadth, and historical demographic reconstruction revealed repeated range contractions and expansions during the Pleistocene, mirroring those observed in macro-organisms. These findings extend the GLQ paradigm to protists, highlighting shared historical and ecological processes across the eukaryotic tree of life and contributing to a unified theory of biogeographic responses to climatic change.

## Introduction

Species geographic distributions and diversification patterns are typically shaped by non-stochastic processes, most notably historical events that influence major climatic changes in the global biota^1^. Among the best-characterised of these events are the Quaternary glacial cycles (between 2.6 Ma and 11.7 ka before present), which have profoundly influenced the distribution and genetic spatial structuring of animals and plants. This succession of warm and cold periods left a trace widely referred to as the “genetic legacy of the Quaternary ice ages” (GLQ hereafter^2–5^). While the effects of the GLQ on the contemporary biogeography of many macroscopic taxa have been widely studied, comparable knowledge is lacking for microbial sized organisms. This gap is particularly evident when considering protists, eukaryotic microorganisms that are neither animals, nor fungi, nor plants but make up the largest part of eukaryotic diversity^6^. Consequently, our limited understanding of protistan biogeographic patterns in an evolutionary framework hinders both the extension of the GLQ paradigm to Domain Eukarya as a whole, which would allow a generalized framework for biogeographic hypotheses grounded on global historical events.

During the Quaternary, a series of ice ages drove the global progression of polar ice sheets and associated biomes toward the equator. Nevertheless, local topography or the proximity of water warm water masses left climatic refugia, geographical zones with milder temperatures, reduced ice cover, or higher water availability, that allowed local populations to survive^2^. Outside these refugia, populations were extirpated, or at least drastically reduced, undergoing genetic bottlenecks. Consequently, refugia acted as biogeographic “islands” where the isolation of incipient lineages promoted diversification. During milder climatic episodes, refugia behaved as sources for the recolonization of surrounding territories. In Europe, the three Mediterranean Peninsulas (i.e. Iberian, Italic and Balkanic) acted as glacial refugia, and were the starting point for northward expansion routes, which eventually produced secondary contact zones between closely related taxa^4,5^. These successive glacial cycles left a lasting imprint on the present-day inter-and intraspecific genetic diversity of many organisms, according to the ecological niche, and the dispersal ability of each species. Among the three major refuge zones, the Iberian Peninsula stands out for its pronounced topographic and ecological climatic heterogeneity. As such, it harbours multiple, internally isolated refugia (appropriately called “refugia within refugia”), generating complex patterns of biodiversity and secondary-contact zones^7,8^. The extensive comparative data available for other macro-organisms, coupled with the high density of distinct internal refugia within a relatively small area (∼580,000 km²), make the Iberian Peninsula an outstanding natural setting for testing the GLQ paradigm in protists.

In order to test the GLQ paradigm, terrestrial species are often preferred over freshwater ones, firstly because river systems and lake basins often have complex and independent geological histories; furthermore, freshwater species are often constrained by water basin^9^. Within protists, however, diversity data on terrestrial assemblages are still scarce, leaving only a handful of taxa with sufficient data to evaluate specific historical hypotheses^10^. One of the best-documented terrestrial protist groups is the order Arcellinida (lobose testate amoebae). This group of protists has been central in a debate on putative microbial cosmopolitanism since the turn of the 21^st^ century^11^. A more recent phylogeographic study has suggested a probable impact of glacial cycles on the distribution of a species complex in the Holarctic realm^12^. Furthermore, their systematics and taxonomy are relatively well resolved^13,14^, and group-specific metabarcoding protocols are available^15^. Together, these features make Arcellinida an excellent model for testing ecological and biogeographic hypotheses, including the GLQ, within continental environments.

Accordingly, our goal was to test the GLQ paradigm on terrestrial protists, using the diversity of Arcellinida in the Iberian Peninsula as a model. We generated *de novo* soil metabarcoding data from across the Peninsula and merged them with every Arcellinida metabarcoding dataset published to date into a single, standardised database, ArKOI. This comprehensive resource allowed us to (i) assess how infra- and interspecific genetic diversity is spatially structured across the Peninsula, (ii) delineate the climatic niches of individual Arcellinida species in Mediterranean and Temperate ecoregions, and (iii) place these patterns in a temporal context. We hypothesize that, despite of their small size, Arcellinida diversity distribution follows similar patterns as macro-organisms. Concretely, intraspecific diversity should be concentrated in the climatic refugia that were already recognised for macro-organisms. Following the GLQ paradigm^2^, Arcellinida populations underwent demographic changes during the Pleistocene. If true, these conclusions would expand the validity of the GLQ to the microbial realm towards a generalized biogeographical theory.

## Results

### ArKOI database

As an initial step toward testing the “genetic legacy of the Quaternary ice ages” (GLQ hereafter^2,3^) biogeographical hypotheses in the Iberian Peninsula^7^, we assembled a comprehensive database of metabarcoding studies targeting Arcellinida. Six published metabarcoding datasets were retrieved (supplementary data S1) and re-assembled with a standardized pipeline (see “Materials and Methods”). We then added *de novo* metabarcoding data from terrestrial samples from locations previously identified as Pleistocene climatic refugia for animals and plants in the Iberian Peninsula, as well as from areas with no documented refugial status^7,8^. The resulting database, called ArKOI (supplementary data S2), contains 615 samples, with 17,023 Arcellinida amplicon sequence variants (ASVs) supported by 26,717,017 reads, clustered into 5,398 operational taxonomic units (OTUs) (supplementary data S3). These OTUs were built following a barcoding gap set in previous studies, and constitute therefore proxies for species in Arcellinida^16–18;^ ASVs correspond to infraspecific diversity, that is, mitochondrial haplotypes (Fig. 1a). ArKOI spans freshwater, soil and saline continental ecosystems across Europe, the Americas and the Middle East. Within the Iberian Peninsula alone, 274 samples produced 2,830 Arcellinida ASVs (1,394,795 reads) clustered into 1,217 OTUs (Fig. 1b). Each sample was assigned to an ecoregion^19^, in the case of the Iberian Peninsula the samples were established in the following ecoregions: 1) Mediterranean Forests, Woodlands (Mediterranean, for short); and 2) Temperate Broadleaf & Mixed Forests (Temperate). This standardized database provides a robust framework for testing biogeographical, ecological and evolutionary hypotheses in Arcellinida.

**Figure 1.**
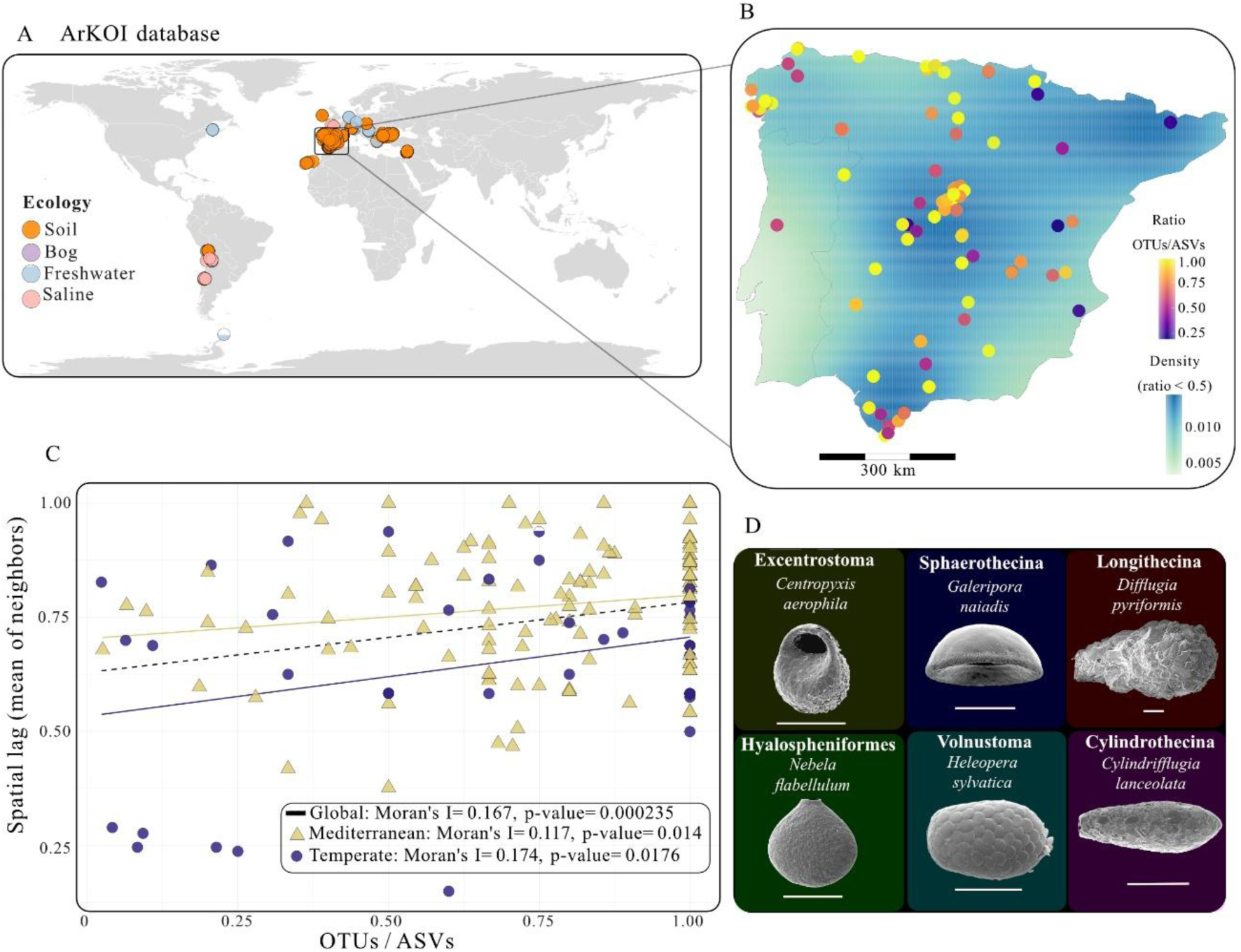
Global coverage of ArKOI and spatial autocorrelation of OTU/ASV ratios in terrestrial Iberian soils. (A) ArKOI database: Global map showing every sample currently included in the ArKOI database, colour-coded by ecosystem type. (B) Iberian Peninsula soil points (KDE samples with ratio < 0.5): Terrestrial samples from the Iberian Peninsula coloured by their OTU/ASV ratio; a kernel-density estimate (KDE) surface highlights diversity hotspots based on samples with ratios ≥ 0.05. (C) Spatial lag vs. Ratio by ecoregions in the Iberian Peninsula: Moran’s I values for OTU/ASV ratios calculated for all Iberian terrestrial samples combined, and each ecoregion analysed separately. (D) Representative scanning-electron micrographs of the major Arcellinida morphotypes; images modified from^14^. Scale bars = 50 µm.

### Spatial autocorrelation of the sample-level Arcellinida ASV/OTU ratio

We determined next how far sites with higher intra-OTU diversity, used here as a proxy for intra-specific diversity, are geographically clustered within the Iberian Peninsula. For each sample we calculated the ASV/OTU ratio of Arcellinida^17^, higher values reflecting greater intra-OTU (equivalent to interspecific) diversity in a given site. Then, we assessed the spatial structure of this ratio with Moran’s Index. Across all Iberian samples we detected a significant, positive spatial autocorrelation (Moran’s I = 0.1668, p = 0.00023; Monte-Carlo observed = 0.16684, p = 0.001), indicating that sites with similar ASV/OTU ratio values tend to be geographically close to each other (Fig. 1b, c). This pattern persisted when the analysis was repeated separately for the two Iberian ecoregions. Both the Mediterranean (Moran’s I = 0.117, p = 0.014) and the Temperate ecoregion (Moran’s I = 0.174, p = 0.0176) showed significant positive autocorrelation (Fig. 1c). The consistent clustering of samples with elevated intra-OTU molecular diversity, observed for the Peninsula as a whole and within each ecoregion, supports the existence of biodiversity hotspots in the Peninsula, a result that is compatible with the existence of Pleistocene climatic refugia as postulated in the GLQ paradigm.

### Informative OTUs from terrestrial ecosystems in the Iberian Peninsula

In order to delimit accurately the location of the biodiversity hotspots, we restricted the analyses to a selection of “informative OTUs” that were abundant and widespread enough to draw conclusions on their evolutionary history at the population level. Following^20^, we classified as “informative OTUs” those OTUs that: (i) contained at least four ASVs, (ii) harboured ≥ 1 % pairwise genetic divergence between any two haplotypes, and (iii) was detected in a minimum of two sampling localities. Although this filter removes OTUs with very low intra-OTU variation (e.g. those affected by recent bottlenecks), it minimises the possible occurrence of sequencing artefacts and produces taxonomic units suitable for population genetics analysis. Applying the criteria defined in the Materials and Methods section, we recovered 341 informative OTUs across the full ArKOI dataset (supplementary data S4).

Next, we identified which of these 341 informative OTUs showed the highest haplotypic diversity in terrestrial ecosystems within the Iberian Peninsula. ArKOI includes sequences from sites situated in several continents and habitat types, which allows identifying possible centres of diversity for each informative OTU. We used the Population-level Haplotype Diversity metric (PH_d_)^20^ to rank OTUs by haplotype diversity; this metric integrates intra-population diversity and the evenness of haplotype distribution across populations. PH_d_ analyses were run separately by ecoregion and by geographical unit (countries). This approach identified 32 OTUs whose highest PH_d_ values occurred in terrestrial systems of the Iberian Peninsula. Of the 32 informative OTUs, 26 showed their diversity peak in the Mediterranean ecoregion, whereas 6 peaked in the Temperate. For both ecoregions, individual informative OTUs displayed median PH_d_ values close to zero, indicating a single dominant ecoregion that can be reconstructed from haplotype-level data (Fig. 2b). All informative OTUs belonged to Infraorder Excentrostoma, the most diverse clade in terrestrial ecosystems^17^.

**Figure 2.**
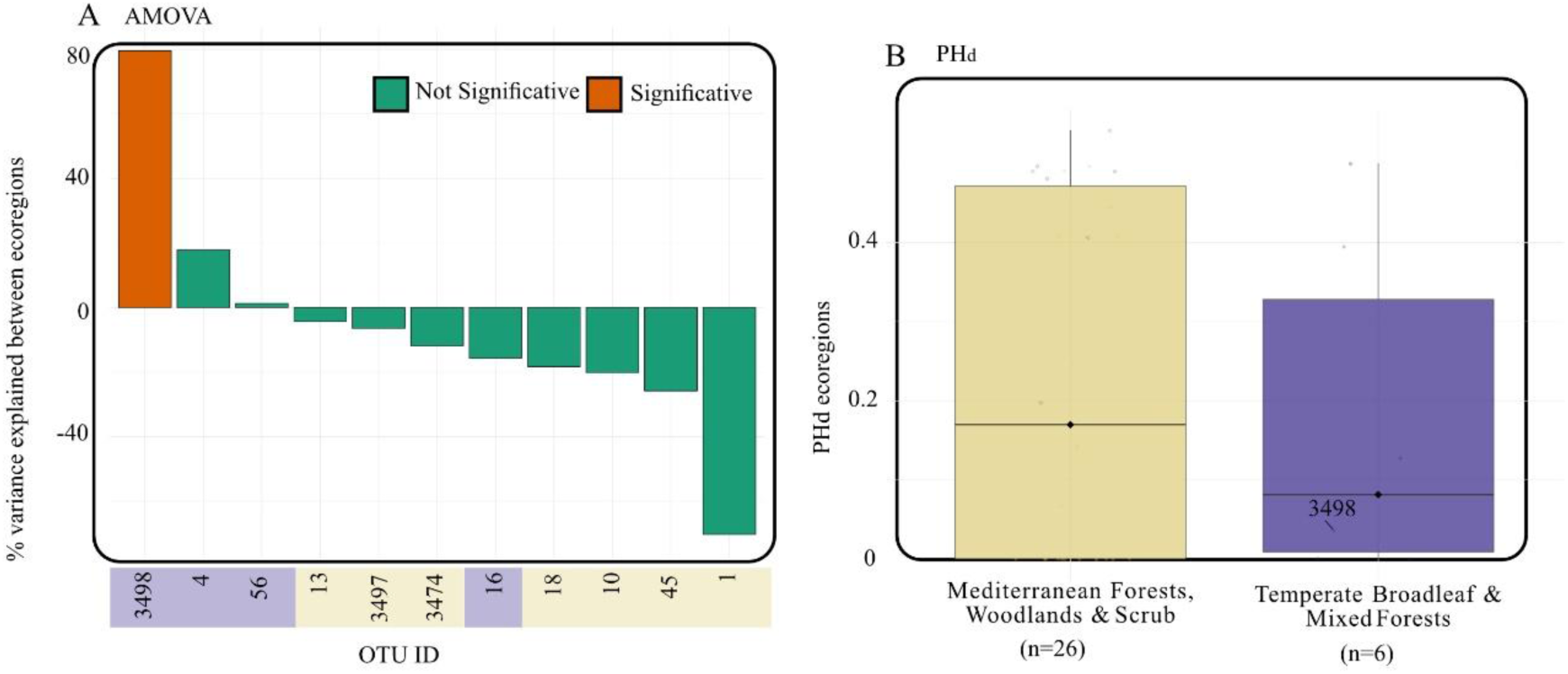
Informative terrestrial OTUs in the Iberian Peninsula. (A) **Percentage of molecular variance explained by ecoregion (AMOVA):** Analysis of molecular variance (AMOVA) for each informative OTU, showing the percentage of molecular variance attributable to ecoregion. (B) **PH_d_ of each OTU per ecoregion:** Results of the PH_d_ analysis for the informative OTUs, summarised by ecoregion.

To test the robustness of these ecoregion assignments in the 32 terrestrial Iberian informative OTUs, we performed an analysis of molecular variance (AMOVA) on Iberian data only. Eleven OTUs (4 Temperate, 7 Mediterranean) occurred in both ecoregions (Fig. 2a). Of these, only OTU3498 showed significant population structure, with 79.52 % of its genetic variance partitioned between ecoregions; but considering all the data outside the Iberian Peninsula it showed a very low PH_d_ (Fig. 2b), which assigns its classification to the temperate ecoregion. The remaining OTUs exhibited low or negative among-region variance, corroborating the PH_d_-based ecoregion classifications in the Iberian Peninsula. We showed therefore, with two approaches, that the 32 identified informative OTUs have their maximum molecular diversity within terrestrial biotopes in the Iberian Peninsula, either in the Temperate or in the Mediterranean ecoregion.

### Geographical spatial structuring of informative species across the Iberian Peninsula

Next, we examined how the genetic diversity of these 32 informative OTUs is distributed geographically across both ecoregions in the Iberian Peninsula. First, we mapped haplotype richness and the proportion of haplotype overlap for every Iberian sample. The Mediterranean informative OTUs show their highest haplotype richness in three areas: the “Sistema Central”, the Baetic region, and the Iberian System (Fig. 3a, S1 and S2). The Temperate informative OTUs display two hotspots of haplotypic diversity in the north-western (Galaic-Portuguese) and north-eastern (Catalonian Pyrenees) corners of the peninsula (Fig. 3a, S1 and S2). Most localities sampled in the Iberian Peninsula contained haplotypes belonging only to OTUs from the same ecoregion; the coexistence between Mediterranean and Temperate OTUs is rare and occurred only in 17 out of the 274 surveyed sites (Fig. 3b).

**Figure 3.**
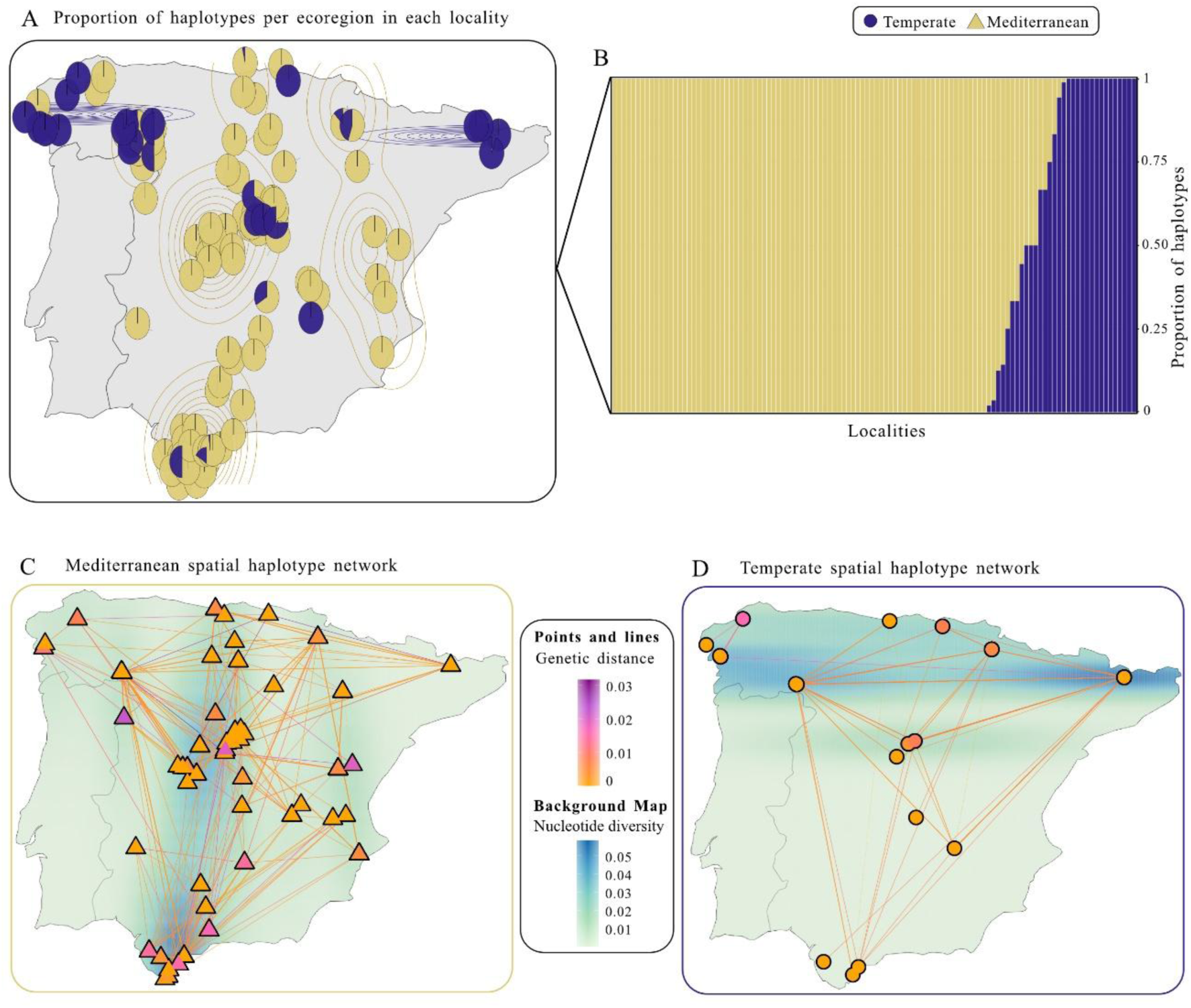
Intra-OTU molecular structuring across the Iberian Peninsula. (A) Relative frequency of haplotypes per ecoregion in samples containing informative OTUs; smooth curves depict kernel-density estimates of the total number of haplotypes per ecoregion. (B) Boxplots showing the proportion of haplotypes per ecoregion for each locality that harbours informative OTUs. (C) For every Mediterranean informative OTU, maps plot haplotype composition by Iberian locality, with point colour indicating nucleotide diversity among haplotypes of each OTU in that locality; connecting lines follow OTU-specific haplotype networks, where colour reflects pairwise genetic distance and dashed lines mark identical haplotypes shared by multiple localities; background colour of the map represents the kde of localities with pi >= 0.005 (D) Same representation as in (C) for informative OTUs from the Temperate ecoregion.

We used an independent approach to corroborate the observed intra-OTU spatial patterns. In that purpose, we combined haplotype-network topology with per-sample nucleotide diversity. The resulting patterns mirrored those based on haplotype richness, with the single exception of the diversity hotspot found in the Iberian System, which was no longer detected. Still, we observed two clear clusters of molecular diversity emerging among the Mediterranean OTUs located in the Central System and Baetic region, respectively (Fig. 3c and S1). The haplotypes found in these clusters were connected to others from surrounding localities by low genetic distances. Temperate OTUs retained the same two northern clusters (Galaic-Portuguese and Catalonian Pyrenees), with peripheral haplotypes likewise linked by low molecular divergence (Fig. 3d and S2). Together, these spatial patterns of intra-OTU diversity matched the locations of well characterized Pleistocene climatic refugia previously identified for animals and plants on the Iberian Peninsula^7,8^. This reinforces the view that Arcellinida experienced range expansion-contractions dynamics around refugia similar to other organisms, in accordance with a generalized GLQ paradigm.

### Climatic-niche structuring of terrestrial informative OTUs across the Iberian Peninsula

Having identified intra-OTU molecular diversity hotspots for the 32 Iberian informative OTUs, we aimed at identifying which present climatic factors explain their present day spatial distribution at best. In particular, we sought which climatic factors might have acted differently on the Mediterranean and Temperate informative OTUs in the Mediterranean and Temperate ecoregions, respectively^21^, in order to better characterize their phylogeographic history. We ran a principal-component analysis (PCA) on the 19 bioclimatic variables plus elevation provided by WorldClim^21^ for every informative OTU. The ordination separated clearly Mediterranean and Temperate OTUs (Fig. 4a). Precipitation-related variables dominate PC1 (Fig. 4b). Furthermore, a one-way Analysis of variance (ANOVA) performed on each abiotic variable (Fig. S3) revealed that factors related to precipitation were the main climatic drivers OTU niches for both ecoregions, and temperature for Mediterranean ecoregion. Such a result is in line with previous works that emphasize humidity as one of the most influential variables in shaping Arcellinida niches^12,16^.

**Figure 4.**
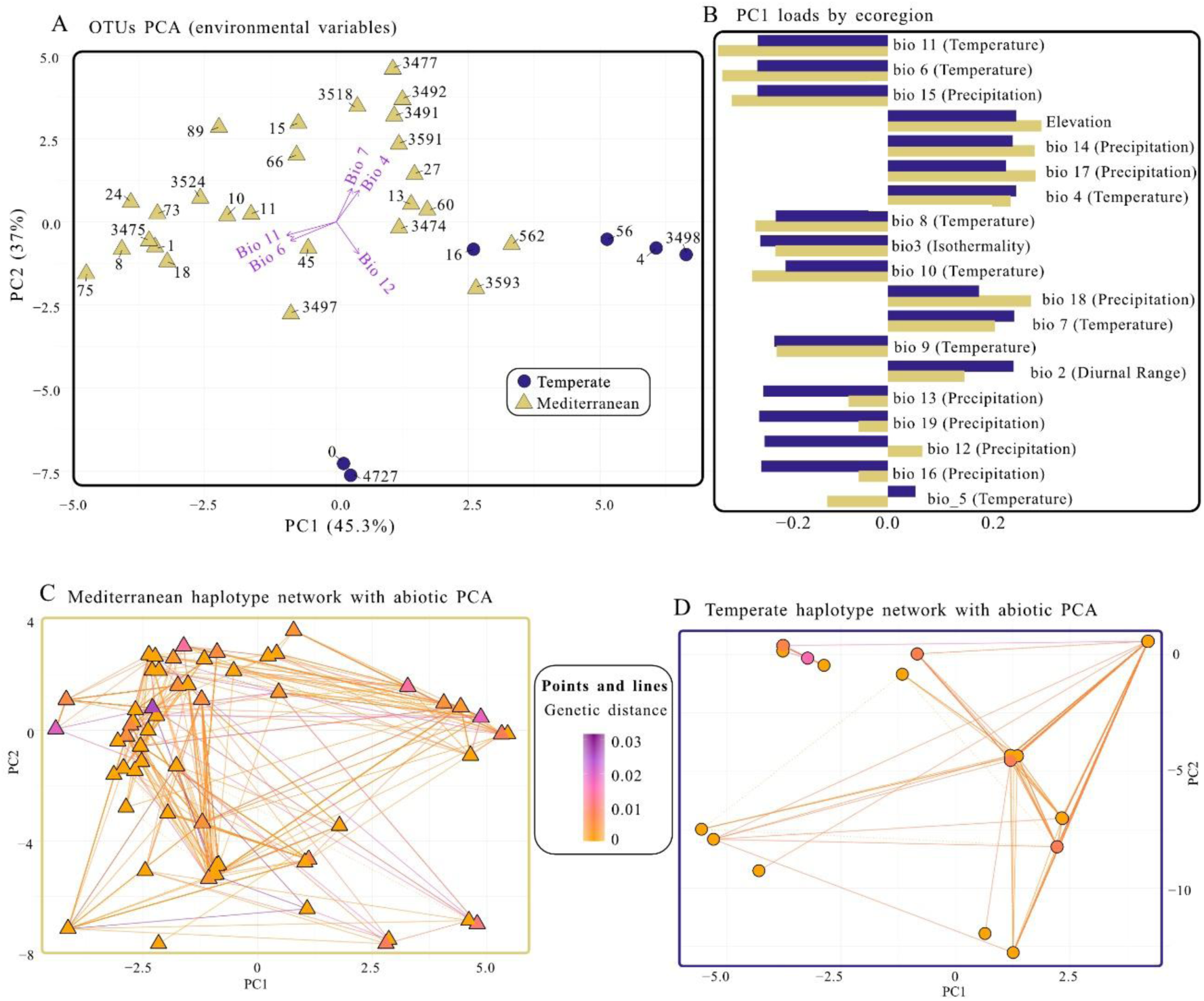
Climatic niche characterisation of Iberian informative OTUs by ecoregion. (A) Principal-components analysis (PCA) of all informative OTUs based on 19 bioclimatic variables plus elevation; points are coloured by ecoregion and the five longest loading vectors highlight the most influential variables. (B) Variable loadings on PC1, shown separately for Mediterranean and Temperate informative OTUs. (C) For each Mediterranean informative OTU, a niche-space PCA plots localities as points coloured by nucleotide diversity among that OTU’s haplotypes at the respective site; connecting lines reproduce the OTU-specific haplotype network, with line colour proportional to pairwise genetic distance and dashed lines indicating identical haplotypes shared by multiple localities. (D) Same visualisation as in (C) for informative OTUs from the Temperate ecoregion.

We obtained a more detailed picture of the distribution of intra-OTU diversity by combining haplotype-network topology with per-sample nucleotide diversity onto the same climatic PCA. Mediterranean OTUs occupy a broad climatic space (i.e. collectively broader niches; Fig. 4c). Most haplotypes fell into one or two dense clusters that harboured the bulk of each species haplotypic diversity; peripheral haplotypes were connected with low genetic distances from these centres and contained little additional nucleotide diversity. In contrast, Temperate OTUs had a noticeably narrower climatic space than Mediterranean OTUs (Fig. 4d). Like for the Mediterranean ecoregion, diversity for each OTU is concentrated into one or two clusters: however, the spread of peripheral haplotypes is noticeably smaller, which mirrors the more restricted Temperate ecoregion climate. The informative OTUs from the Mediterranean and Temperate zones displayed clear climatic-niche differentiation, driven primarily by precipitation regimes. These results reinforce the view that climate plays a key role in shaping the spatial structuring of Arcellinida lineages, which do have different climatic optima. These preferences shape present and most probably past distribution of Arcellinida. This is consistent with the survival of the species within different Pleistocene climatic refugia.

### Climatic niche overlap between Mediterranean and Temperate informative species

Given the different genetic diversity within Mediterranean and Template OTUs, we hypothesized that both groups should have probably different dispersal histories, which would be best reflected by determining how far the niches of the informative OTUs overlap. As a first step, we compared the niche breadth and mean intra-OTU genetic diversity of the 32 Iberian informative OTUs using Iberian records only. Mediterranean OTUs showed markedly larger median niche hypervolumes (43.94) and longer mean intra-OTU distances (0.01149) than their Temperate counterparts (24.30 and 0.0066, respectively; Fig. 5a,b). We tested a possible relationship between geographic area and the number of haplotypes of each OTU per ecoregion by performing a Spearman’s rank correlation analysis. The Spearman rank test revealed a positive correlation between geographical range and haplotype number in Mediterranean OTUs (ρ = 0.47, p = 0.016; Fig. S4). In contrast, such relationship was not significant in Temperate OTUs (ρ = -0.14, p = 0.80; Fig. S4), suggesting that Temperate lineages experienced a range-restricting bottleneck.

**Figure 5.**
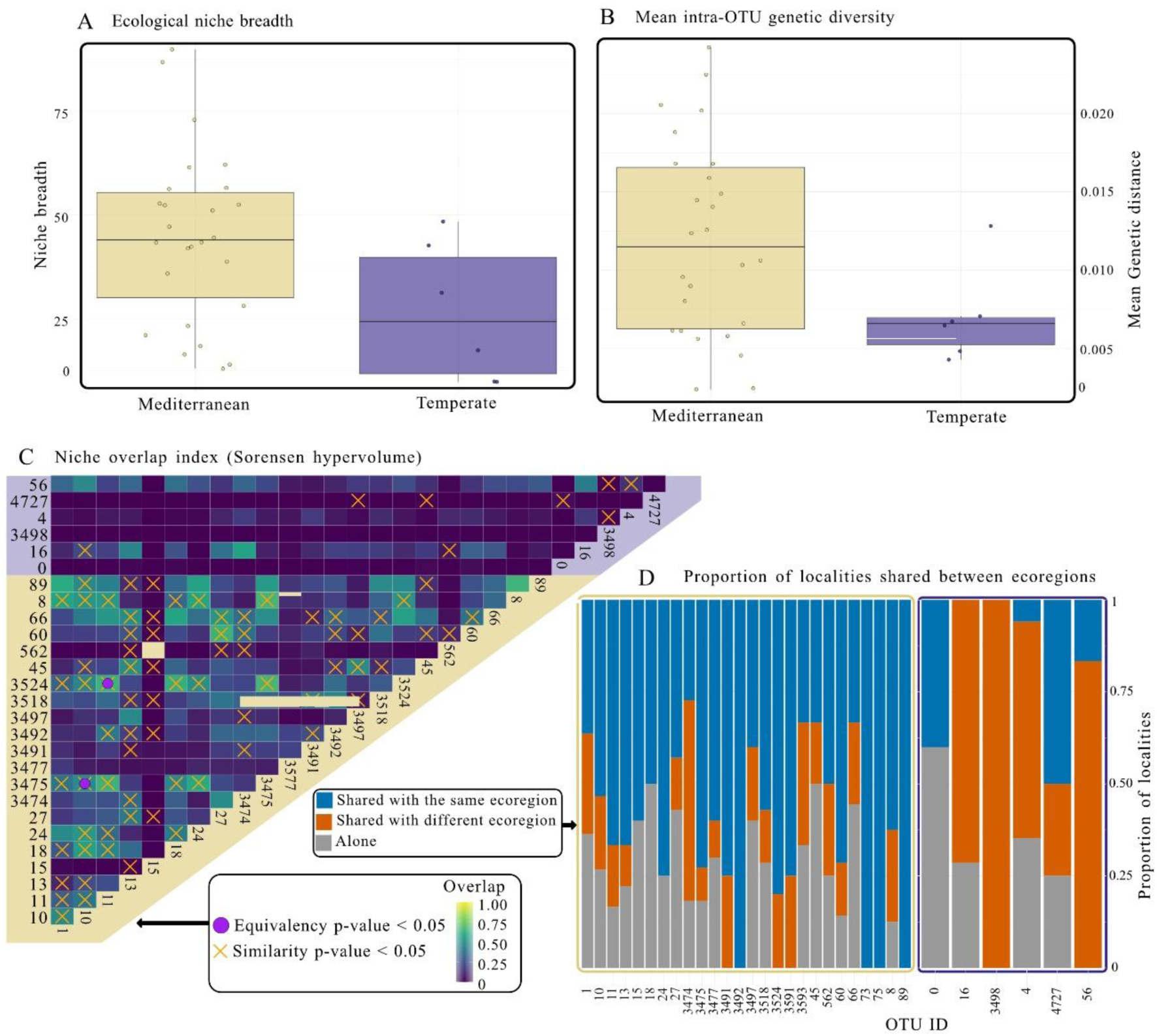
Climatic-niche and geographic overlap among Iberian informative OTUs by ecoregion. (A) plot of niche breadth for each Iberian informative OTU, grouped by ecoregion. (B) plot of the mean intra-OTU geographic distance for each Iberian informative OTU, grouped by ecoregion. (C) Pairwise Sørensen indices of climatic-niche overlap based on hypervolumes calculated for each informative OTU; crosses indicate niche similarity significantly higher than expected by chance, whereas filled circles denote full niche equivalence. (D) Boxplot showing the percentage of Iberian localities in which each informative OTU co-occurs with haplotypes of another informative OTU from a different ecoregion, co-occurs with an OTU from the same ecoregion, or occurs alone.

To calculate the degree of climatic niche overlap between OTUs, we generated hypervolumes from PCA axes of the climatic variables described above, which show that OTUs overlap far more within than between ecoregions (Fig. 5c). The strongest cross-ecoregion overlap involved Temperate OTUs OTU16 and OTU56 with Mediterranean hypervolumes. Both niche-equivalence and similarity tests confirmed that most climatic niches are more similar than expected by chance only within the same ecoregion. The only exception consisted in the overlap between Temperate OTUs OTU16 and OTU4727. Finally, the locality-sharing analysis (Fig. 5d) revealed that almost every Temperate OTU coexists with Mediterranean lineages, OTU0 being the only exception. In turn, Mediterranean OTUs co-occur almost exclusively with other Mediterranean OTUs. This asymmetric pattern indicates that Temperate taxa occupy a subset nested within the broader Mediterranean distribution, consistent with a post-glacial northward range expansion that started from Mediterranean refugia as the climate became increasingly warm and dry.

### Temporal context in informative species

A temporal framework is still necessary in order to determine the implication of Pleistocene climate shifts in present-day Arcellinida spatial molecular diversity patterns. Because no standard COI substitution rate exists for Arcellinida, we used as a reference date the splitting of the *Hyalosphenia papilio* “shadow-species” complex, corresponding with the development of the boreal *Sphagnum* peatlands to which it is confined^12^. Assuming the diversification of these amoebae followed the inception of boreal raised bogs (between 7 and 20 Ma ago^22,23^), we dated COI trees under several molecular-clock models. All models converged on a lineage-specific rate of ∼0.014 substitutions site^-^¹ Myr^-^¹ within lineage (Fig. 6a), a value comparable to estimates for most animal phyla^24^ (supplementary data S5; Fig. 6b). Although no truly universal clock exists, this rate provides a useful proxy for placing diversification and demographic events on an approximate timescale in Arcellinida COI.

**Figure 6.**
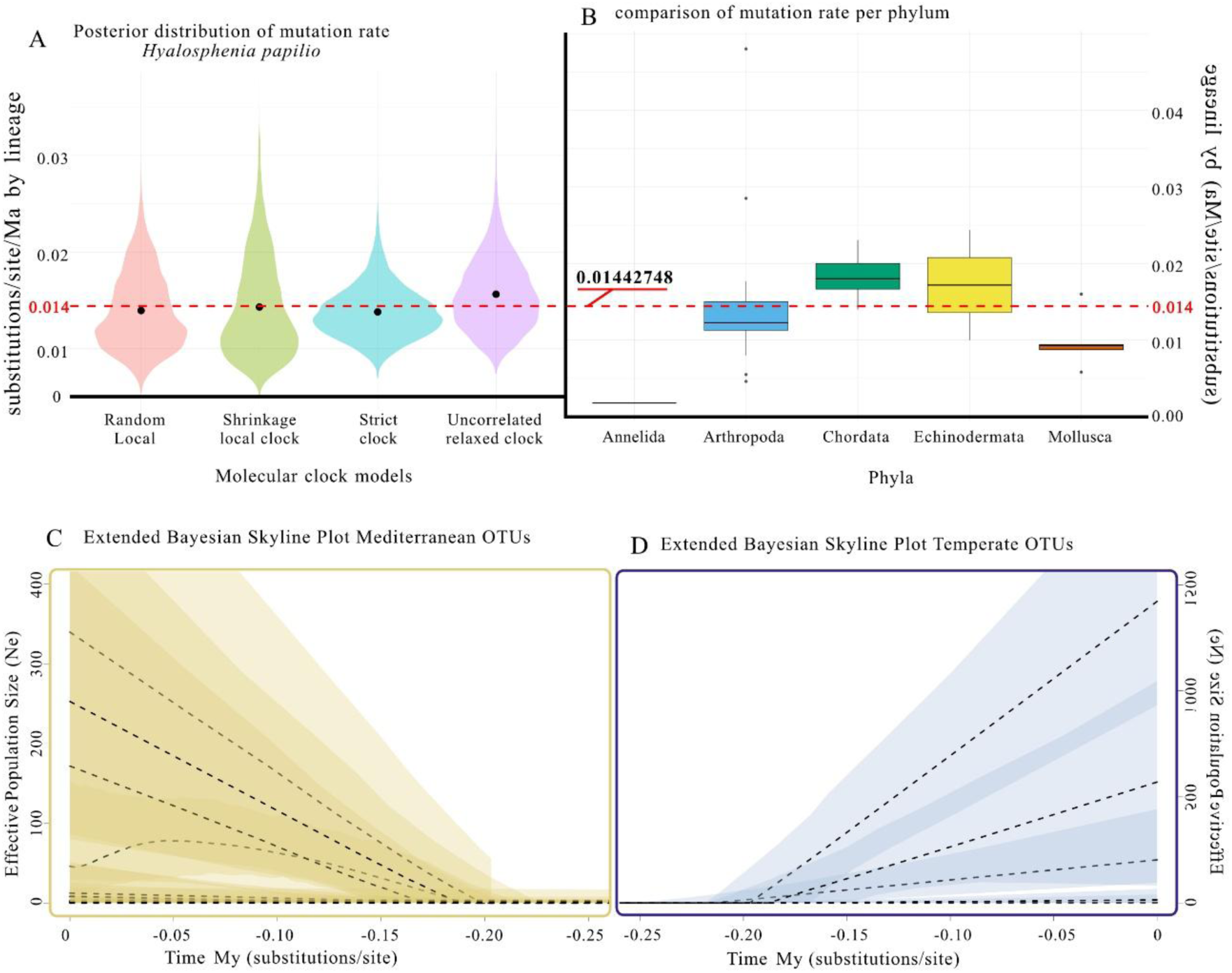
Temporal framework for Arcellinida. (A) Violin plots of branch-specific substitution rates (subs/site⁻¹/Myr⁻¹) estimated under each molecular-clock model; the dashed horizontal line marks the mean across clocks. (B) Comparative box-plot of branch substitution rates for major metazoan phyla, plotted alongside the mean Arcellinida rate (0.014). (C) Extended Bayesian Skyline Plot (EBSP) showing demographic history of Mediterranean informative OTUs: dashed lines, median effective population size; shaded envelopes, 95 % highest-posterior density. (D) Same EBSP representation for informative OTUs from the Temperate ecoregion.

Using it in Extended Bayesian Skyline Plot analyses under a strict clock, we reconstructed effective population-size (Ne) trajectories for each OTU. Where resolution permitted, population shifts cluster around 200 ka, well within the Pleistocene, despite the limitations imposed by an inferred rate and the short (∼360 bp) metabarcoding ASV fragments. Intriguingly, Temperate OTUs show larger Ne than Mediterranean OTUs, despite their lower haplotypic diversity and narrower climatic niche. This suggests that both lineage groups persisted through the Pleistocene but experienced different demographic dynamics.

## Discussion

### Geographical structuring of potential Iberian refuges for Arellinida

Our results indicate that the genetic structuring of the different Arcellinida OTUs aligns with Pleistocene climatic refugia previously characterized for multicellular plants and animals. This demonstrates that the GLQ paradigm can also apply to at least some unicellular microorganisms. Furthermore, our findings support the “refugia within refugia” hypothesis that states that the Iberian Peninsula, which constitutes a refuge as a whole, includes distinct, independent centres of high intraspecific molecular diversity^7,25^. We identified potential climatic refugia for Temperate and Mediterranean ecoregions, respectively.

For the Temperate ecoregion, we identified two main areas of high intra-OTU diversity: the Galaic-Portuguese region in the west and the Eastern Pyrenean region in the east (Fig. 3d and S2). The Galaic-Portuguese region acted as a refuge for taxa that expanded their distribution during the Holocene, such as, for instance, the Iberian frog *Rana iberica*^26^ or the pedunculated oak *Quercus robur*^25^. This area likely acted as a climatic refuge during the Quaternary due to several factors: i) the Atlantic Ocean acted as a thermal buffer, moderating extreme cold during glaciations; ii) the Atlantic influence also brought high levels of humidity; and iii) the mountainous terrain (e.g., “Sierra de Ancares”, “Macizo Galaico”) protected the region from cold winds. Similarly, the Catalonian Pyrenees region has been recognized as a refuge for various plant and animal species, such as the viviparous lizard *Zootoca vivipara*^27^ and the downy oak *Quercus pubescens*^25^. Here, the moderating influence of the ocean, and the complex topography, with valleys that were sheltered from north winds preserved deciduous and mixed forests (e.g., beech, oak) during glaciations.

For the Mediterranean ecoregion, we identified two areas of high intra-OTU diversity that likely acted as Pleistocene climatic refugia for Arcellinida: the “Sistema Central” (Central system), corresponding to the Guadarrama and Gredos mountain ranges in the centre of the Peninsula, and the Baetic Range, in southern Iberia (Fig 3c and S1). The Central Plateau of the Iberian Peninsula has been described as a climatic refuge for various plant and animal species, such as the Iberian green lizard *Lacerta schreiberi*^28^ or the midwife toads of genus *Alytes*^29^. This area is characterized by its mountainous zones, including the “Sistema Central” (e.g., “Sierra de Guadarrama”, “Gredos”) and the “Montes de Toledo”, whose internal valleys preserved milder and wetter microclimates^30^. However, the extent of these refugia was more limited and fragmented compared to the north of the Iberian Peninsula. The continental position (far from the sea) and the high elevation of the Central system made it less suitable as a large-scale refuge. Therefore, the central Iberian Peninsula most likely resembled a mosaic of microrefugia rather than a broad, continuous refugial area. In contrast, the southern Iberian region has been identified as a refugium for various species, such as *Forficula aeolica*^31^, *Timon lepidus*^32^, and the microscopic rotifer *Brachionus manjavacas*^33^ amongst others. This region experienced a relatively mild climate, especially in coastal and lowland areas, which remained largely ice-free. The Mediterranean Sea and Atlantic Ocean acted as climatic buffers, reducing temperature extremes, while the sheltered valleys and shaded areas of the Baetic region kept higher humidity and milder temperatures.

It is important to note that not all refugia were equally suitable during each glaciation, and not all refugia were suitable for all species. Even though some populations began to diverge before the Pleistocene, their continued presence in the same areas could suggest that these regions remained suitable for survival during the ice ages in the Pleistocene. Finally, it is important to emphasize that many more refugia have been characterized in the Iberian Peninsula than those recovered in our study^7,25^, and that a more complete sampling would probably reveal more haplotypic diversity hotspots.

### Sanctuaries vs. climatic refuges in the Iberian Peninsula for Arcellinida

By integrating haplotypic diversity and genetic distance data within each informative OTU independently, we were able to identify geographic areas with higher diversity and relate them to known climatic refugia. Furthermore, Pleistocene phylogeographic diversity patterns can provide information on how ice age episodes influenced the different taxa. Although phylogeographic patterns depend on each taxon life history and are therefore known to be species-specific, they can be broadly classified into two models^34^: i) the Sanctuary model (S model), where Pleistocene glaciations likely caused population fragmentation and promoted lineage sorting, but allowed the preservation of, at least, part of the ancestral diversity accumulated before the glaciations; and ii) the Refuge model (R model), where only a small portion of the ancestral population survived in the climatic refugia, and most of the ancestral intraspecific diversity was lost. In the S-model, larger genetic distances can be expected, given the fact that populations had diversified before glaciations. In turn, in the R-model, a much shallower mitochondrial diversification is expected due to populations bottlenecks associated with founder effects caused by the effects of glaciations themselves. Furthermore, if different populations have remained isolated in separate refugia, the geographic contact zones where these lineages meet during post-glacial expansion are expected to exhibit the highest genetic diversity compared to the original refugial populations^35^.

Our results for Arcellinida broadly demonstrate that most OTUs associated with the Temperate ecoregion fall within the refugium model (R model). When applying a threshold of >1% mean intra-OTU divergence to classify OTUs according to refugial models (Fig. 5b), we found that, among the Temperate ecoregion OTUs, only one out of six meets the criteria for the Sanctuary model (S model). In contrast, 15 out of 26 Mediterranean OTUs would fall within the S model (Fig. 5b). Thus, OTUs from the Temperate ecoregion exhibit very low average molecular diversity (Fig. 5b), both in terms of haplotypic richness and genetic distances, indicating a high degree of homogeneity. The temperate zone of the Iberian Peninsula is primarily a colonization area for species from temperate Europe, representing the southern range limits of many taxa. These species may have had their centre of diversity in higher latitudes but these populations may have been extirpated. This peripheral distribution likely explains the dominance of Refugium model (R model) patterns in this ecoregion in the Iberian Peninsula. On the other hand, the majority of Mediterranean ecoregion OTUs can be classified within the Sanctuary model (S model), showing much higher average molecular diversity compared to the Temperate OTUs. These OTUs also exhibit a more heterogeneous pattern of genetic structuring across the Iberian Peninsula, with distinct centres of higher molecular diversity corresponding to the climatic refugia. This demonstrates that the neutral genetic diversity of populations is more strongly shaped by their position relative to historical refugia than by the current core of their distribution range.

### Arcellinida climate niche facing the future challenge of climate change

Our results confirm that wide scale climate changes play a major role in shaping the geographical structuring of intraspecific diversity in Arcellinida, following mechanisms that differ depending on the ecoregion. Precipitations is the primary factor shaping the current geographic distribution of OTUs from the temperate ecoregion (Fig. 4a, b and S3). Previous studies have demonstrated that humidity is one of the most important variables in the biogeography of protists in general^10,36^. In Arcellinida, depth to water table (a proxy for humidity) is one of the most relevant diversity drivers in peatlands; moreover, transitions between aquatic and terrestrial ecosystems are rare, having occurred only a few times in their evolutionary history^16^. In contrast, temperature-related variables were also important drivers for those OTUs associated with the Mediterranean ecoregion. This suggests that the distribution of Arcellinida OTUs is shaped not only by fine-scale microhabitats, but also by broader climatic conditions, implying that the ongoing global warming could profoundly impact the distribution and population dynamics of these species.

One of the greatest challenges of the Anthropocene is climate change, marked by rising global temperatures^37^. The Iberian Peninsula is projected to be one of the most strongly affected regions in Europe, with significant temperature increases and precipitation declines leading to progressive aridification^38^. Under these conditions, there is evidence of a northward migration of (macroscopic) species into higher latitudes within very short evolutionary timescales^39,40^. Conversely, species adapted to temperate climates are experiencing population declines, in effective size, ecological function, and geographic distribution^40,41^, placing them at risk of extinction^42^. The distribution patterns observed for Arcellinida OTUs mirror these trends described for macroorganisms. Temperate ecoregion OTUs exhibit more restricted distributions and narrower niche breadths, compared to Mediterranean ecoregion OTUs (Fig. 5a and b). In contrast, Mediterranean OTUs display wider geographic distributions and broader niche amplitudes. These Mediterranean OTUs are also found, although in relatively low proportions, in the northern Iberian Peninsula (within the temperate zone). They present low nucleotide diversity in northern localities, and have close genetic similarity to southern populations (see previous discussion) which suggests that their presence is likely the result of recent colonization events. Thus, our results indicate that Mediterranean Arcellinida OTUs are in a process of expansion towards the northern, temperate regions of the Iberian Peninsula, consistent with broader patterns of climate-driven range shifts.

### The importance of temporal context in biogeographical inferences

Current distribution patterns are the result of stochastic processes and historical events that have shaped the evolutionary history of species^43^. Therefore, to infer causal drivers and ecological mechanisms underlying present-day distribution patterns and community structures, it is essential to incorporate a temporal framework that allows correlating these patterns with historical events. Such information can usually be obtained by estimating divergence times using fossil records and molecular clocks^13,44,45^. In the case of Arcellinida, inferring accurate evolutionary timelines is particularly challenging due to the difficulty of assigning fossils to taxa. Indeed because of inconspicuous morphologies, evolutionary convergences, phenotypic plasticity or cryptic diversity^14^ are common in Arcellinida, fossil records are at best fragmentary. To overcome these limitations, an indirect calibration using known cladogenesis events has to be applied for calibrating molecular clocks. Here, we used the diversification of the *Hyalosphenia papilio* species complex in peat bogs, driven by the habitat specificity of these species for *Sphagnum* dominated peatlands^12^, which provided a basis for calibrating a molecular clock for the COI marker in Arcellinida. While there is no universal molecular clock applicable to all genes, studies on COI in Metazoa have reported relatively consistent substitution rates across lineages, typically within a range of ±0.01 substitutions per site per million years within lineages (not between lineages), except for specific phyla or taxonomic groups (Fig. 6b). Our estimated substitution rates for the COI molecular clock in Arcellinida fall within these established ranges for Metazoa (Fig. 6a and b).

Our results, based on this molecular clock calibration, indicate changes in effective population sizes between approximately 150,000 and 200,000 years ago (Fig. 6 c and d). This corresponds with Pleistocene climate cycles, particularly the transition from the interglacial Marine Isotope Stage MIS 7 to the glacial MIS 6^46,47^. The intensification of glacial cycles during the Mid-Pleistocene Transition (MPT), marked by longer and more severe glacial periods, likely had a profound impact on habitat availability, resource distribution, and the connectivity of refugial populations. These patterns mirror the well-documented refugia model described for plants and animals in Europe during the Quaternary^2,3,48^. By situating our demographic inferences within the well-established timeline of Quaternary glacial-interglacial cycles^2,48^, we can confirm the implication of QLC in Arcellinida, and possibly other protists groups. Such results suggest a generalization of QLC to the Domain Eukarya as a whole, no matter the size of the organisms.

## Materials and methods

### eDNA extraction, amplification and sequencing

118 Soil samples were collected across Iberian Peninsula (Spain and Portugal) and Europe (France, Ireland, Czech Republic and Switzerland) (supplementary data S1), in order to correctly characterise the geographical centres of origin of the different operational taxonomic units (OTUs). Environmental DNA (eDNA) extraction, amplification and sequencing were performed as described in previous studies^15^, and followed the recommendations^49^. Total eDNA was extracted using the DNeasy PowerSoil Pro Kit (Qiagen) following the instructions provided by the manufacturer and preserved frozen (−20°C) until further processing. For each round of eDNA extraction, a negative control (extraction without sample) was performed to track potential contaminations.

We amplified a portion of 640 bp from the cytochrome oxidase subunit I (COI), using a two-step nested polymerase chain reaction (PCR) protocol specifically designed for Arcellinida. For the first PCR, we used the universal COI primers pair LCO-1490 as forward and HCO-2198 as reverse^50^, and for the second PCR, we used LCO-1490 as the forward primer and the Arcellinida-specific primer ArCOIR as reverse^51^; for this second PCR, we used a unique combination of primers with tags for each sample, as described in^15^.

Finally, we quantified the PCR products using a Qubit 3 fluorometer (Invitrogen), with dsDNA high-sensitivity (HS) assay kits (ThermoFisher), for normalized the DNA concentration. Then, the normalized PCR products were sequenced using a MiSeq 500 cycles v2 paired-end 250 bp. All sequencing was performed in the Genomics Unit of the Fundación Parque Científico de Madrid, Spain.

### ArKoi database

We selected metabarcoding studies specifically targeting Arcellinida that utilized the ArCOI primers (supplementary data S1). For each of these studies, raw sequence data were independently assembled de novo following the standardized protocol and format described in^20^. The reads per sample were processed using the DADA2 package in R^52^. Chimeric sequences were subsequently removed using the *removeBimeraDenovo* function. After obtaining the ASVs and read counts per locality, tag-jumping artefacts were removed using the custom script script_0 (supplementary data). This script first filters sequences based on length, excluding those shorter than 300 nucleotides and longer than 325 nucleotides, as sequences taxonomically assigned to Arcellinida are known to fall within this range^15,17^. Then, we proceeded in minimizing the effects of tag jumping by eliminating potentially artefactual occurrences between localities. Tag jumping is a common artefact in metabarcoding studies, caused by the coalescence of clusters on the Illumina flow cells originating from different template molecules or growing into one another, resulting in false-positive assignments when signal dominance shifts within a single cluster^53^, among other factors^17^. In order to minimize possible false-positive locality assignment we used the same custom script script_0 (supplementary data), where the proportion of reads for each amplicon sequence variant (ASV) in a given locality relative to the total reads of that ASV across all localities were calculated. Based on these proportions, a filtering threshold of 10% was applied: ASV occurrences in samples/localities with relative abundances below this threshold were removed from the final dataset. These false-positives occurrences typically belong to dominant ASVs, in terms of total reads, but appear with low reads in samples where they should be absent.

The resulting FASTA files were transformed in the following format: >sequence_ID merged_sample={ ’LOCALITY_ID1’: reads, ’LOCALITY_ID2’: reads, … }, accompanied by a metadata file containing information associated with each locality ID. All localities were assigned to an ecoregion based on [XXX] and abiotic variables derived from WorldClim^21^ (accessed on 19/03/2025), using shapefiles (.shp) from these sources and the latitude/longitude of each sample.

All individual FASTA files retrieved from the different metabarcoding studies were merged into a single file named ArKOI_database.fasta. Each unique ASV was assigned a unique identifier in the format ArKOIX (where X is a unique number), and locality information was standardized and recorded. The final FASTA file was then checked again for chimeras using VSEARCH (v2.14.1)^54^.

Finally, ASV taxonomic assignment was conducted using the eKOI taxonomy database^55^, implemented through a custom script (5_taxonomic_assignation.py). This script generates a separate folder for each FASTA file and creates an Excel file with taxonomic assignments based using VSEARCH. ASVs with less than 84% similarity to reference Arcellinida sequences were excluded as not belonging to Arcellinida (threshold determined empirically based on local phylogenies^15,17^). Subsequently, a FASTA file containing only Arcellinida ASVs was generated, forming the final ArKOI database (supplementary data S2).

### Operational Taxonomic Units (OTUs)

The custom scripts from^20^ was used to retrieve each ASV’s ID and match it with metadata to create a data frame containing the sequence ID, geographic coordinates, reads, ecoregion and climatic data. Because some ASVs were found in multiple localities, duplicates were generated for each locality using the suffix _dupN (where N indicates the duplicate number), ensuring unique identifiers for each ASV-locality combination. To avoid biases due to replicated samples, only one ASV per locality was retained, and duplicate ASV IDs and geographic coordinates were removed. A final FASTA file was generated, integrating ASVs across localities, and aligned using MAFFT (v7.490)^56^.

OTUs were inferred from the resulting alignment using the scripts from^20^. VSEARCH was employed to cluster ASVs at a 97% identity threshold, a value previously shown to a good approximation to delineate Arcellinida species^17,18^. Previous works showed that slight modifications of this threshold did not significantly impact biological interpretations^18^. Representative sequences of each OTU (centroids) were saved in a FASTA file (otus.fasta), and a .uc file (otus.uc) mapping each ASV to its corresponding OTU was also generated. Finally, the .uc file was processed to produce a tab-delimited text file (otus_mapping.txt) recording the assignment of sequence IDs to OTUs.

### OTU/ASV ratio per sample

We evaluated the spatial distribution of the ratio between OTUs and ASVs taxonomically assigned to Arcellinida obtained from terrestrial samples from the Iberian Peninsula. For each locality, the OTU/ASV ratio was calculated following^17^. Localities with a ratio < 0.5 were analysed with a kernel-density estimate (KDE) using the script script_1 (supplementary data). To assess spatial structure, we constructed a k-nearest-neighbours matrix (k = 4) and computed global spatial autocorrelation with Moran’s I, using 999 Monte-Carlo permutations. Moran’s I was also calculated separately for each ecoregion. Finally, we produced a scatterplot of the OTU/ASV ratio versus its spatial lag (mean of the four nearest neighbours), overlaying both a global regression line and ecoregion-specific regressions.

### Selecting informative OTUs

Once the OTUs were obtained, the relationships between genetic and geographic distances of the sequences were analysed. We converted the sequence alignment into a DNAbin object suitable for calculating genetic distance matrices using the K80 model. Additionally, a matrix of geographical distances was calculated using the coordinates of the localities associated with the sequences, employing the *distVincentyEllipsoid* function. To ensure robust biogeographical analyses and minimize potential sequencing artefacts, we filtered the data to retain only informative OTUs, defined as those (i) containing at least four sequences from a minimum of two distinct locations and (ii) containing at least one pair of sequences that differ by ≥ 1% (uncorrected p-distance) following^20^.

### Population structure and ecoregion characterization per informative OTU

Haplotype networks were subsequently constructed for each informative OTU using the haploNet function^57^. These networks, based on Minimum Spanning Networks (MSNs) were constructed and used in subsequent analyses. Then, we examined genetic structuring both among and within ecosystems and ecoregions. To do so, we applied a modified haplotypic-diversity index (H_d_) that yields PH ^20^, which represents the proportion of haplotypes shared among populations. For each informative OTU, the ecoregion contributing the largest relative share of haplotypes was considered as putative source population. We selected the informative OTUs to be kept for further analyses based on a series of filters, as follows: i) Ecosystem filter; we retained only those informative OTUs assigned solely to terrestrial ecosystems based on their PH_d_ reconstruction; and ii) Geographic filter; from this subset, we selected only OTUs recorded in the Iberian Peninsula (Spain and Portugal). Finally we perform an ecoregion assignment (Temperate or Mediterranean), of the terrestrial Iberian informative OTUs, according to their PH_d_ values. Consequently, the set of informative OTUs analysed here consists of soil-dwelling lineages from the Iberian Peninsula that have been assigned to an ecoregion via their PH_d_ profiles.

To ensure that there is no effect of ecoregion on intra-OTU diversity of the informative OTUs that could bias the biogeographic interpretations, we calculated haplotype diversity (Hd) and branch diversity (Bd) using the terrestrial Iberian informative OTUs. These metrics were computed using three different approaches: based on the number of ASVs per haplotype, the total number of reads, and a logarithmic transformation (log(reads+1)). This approach is particularly useful for communities with high variability in ASV abundances/reads, as it allows patterns of dominance or inequality to emerge. Additionally, we calculated nucleotide diversity (π) using the *nuc.div* function^57^ and performed Tajima’s D test using the *tajima.test* function^57^. Given the filtering criteria applied to the informative OTUs based on ASVs numbers and genetic distances, negative Tajima’s D values could be expected. Finally, to compare across ecoregions, we first tested for normality using the Shapiro-Wilk test, which indicated that the data did not meet the assumptions of normality for any variable. Consequently, we employed the non-parametric Kruskal-Wallis test to assess the effect of ecoregion in this metrics. Pairwise comparisons between ecoregions were then conducted using Dunn’s Kruskal-Wallis multiple comparisons test.

### Analysis of molecular variance (AMOVA) per Iberian Peninsula informative OTU

To test whether each informative OTU (defined on the PH_d_ metric) is correctly assigned to an ecoregion, we ran an analysis of molecular variance (AMOVA) with the script script_2 (supplementary data). For every OTU individually, we (i) built a genetic-distance matrix, and (ii) performed a hierarchical analysis of molecular variance (AMOVA) (region / site) with 999 permutations. For each OTU we recorded the number of sequences, sites and regions sampled, the percentage of molecular variance attributable to differences among regions, and the associated p-value.

### Spatial distribution of informative OTUs across the Iberian Peninsula

To illustrate the relative haplotype richness per locality, and geographic overlap between the different ecoregion OTU we used the script script_3_a (supplementary data). In that purpose we mapped the proportion of unique haplotypes from each ecoregion that occur in every sampling site, displaying them as pie charts with the proportion of overlap per ecoregion. Kernel-density contours were added separately to illustrate overall haplotype density per ecoregion We then mapped the haplotype networks per informative OTUs in the Iberian Peninsula map, using the script script_3_c (supplementary data). Nucleotide diversity (π) was calculated per locality for each OTU (with respect of the haplotypes shared in the locality by each informative OTU) and used to colour the haplotypes in the geographic map. A background KDE (Kernel-Density Estimate) were performed with the function kde2d, based on localities with π ≥ 0.005. The haplotype networks of all informative OTUs were covered: haplotype links are coloured by genetic distance, and dotted when a haplotype occurs in multiple localities. The resulting figure simultaneously shows sampling effort, genetic connectivity among haplotypes, and within site molecular diversity.

### Correlation between haplotype richness and geographic range

We tested whether the number of haplotypes per informative OTU was correlated with its geographic range within each ecoregion using the script script_S5 (supplementary data). For each OTU we used the record coordinates to delineate a convex hull and calculated its area (km²) as a proxy for range size. Haplotype richness and range size were then analysed with Spearman’s rank correlation (cor.test in R), separately for Mediterranean and Temperate ecoregions, to determine whether OTUs occupying larger areas harbour more haplotypes. Results were visualised in a log– log scatterplot with points, trend lines, and annotations of ρ and p-value.

### Ecology of the informative OTUs

We characterized the environmental niche of each informative OTU of the Iberian Peninsula using the script script_4_a_b (supplementary data). As a proxy for OTU climate niche, WorldClim bioclimatic variables and altitude were extracted^21^, and centroids (means of all climatic variables) were computed for each informative OTU. These centroids were subjected to a global PCA. We did a one-way ANOVA for every environmental variable identified that differed significantly between the Mediterranean and Temperate ecoregions; variables were ranked by F-statistic in a bar plot. Separate PCAs were then run for each biome, and the PC1 loadings of both analyses were plotted side by side to highlight the main environmental drivers of niche variation within each ecoregion.

We projected the haplotype network into a “climatic space” to integrate the ecological and genetic data, with the script script_4_c_d (supplementary data). First a PCA of the abiotic variables was calculated separately for the haplotypes of each informative OTU per ecoregion. Then, nucleotide diversity (π) for every OTU-locality combination (representing sets of haplotypes sharing the same abiotic conditions, as they share the same locality), as described above, was added to the plot. Haplotype-network links between haplotypes, coloured by genetic distance (dotted when one haplotype occurs in several localities), were covered.

### Comparison of niche breadth and genetic distances by ecoregion

We estimated ecological niche breadth for every informative OTU in each ecoregion with the script script_5_a (supplementary data). For every ASVs belonging to an OTU we extracted the relevant climatic variables, then defined niche breadth as the mean of the standard deviations of those variables; concretely, the amount of environmental variation encompassed by the informative OTU. Mean genetic distances per ecoregion were calculated with the script script_5_b (supplementary data); which computes the average pairwise genetic distance for each informative OTU within each ecoregion. We used a 1% mean intra-OTU divergence threshold to classify OTUs according to the refugial models: Sanctuary (S model) and Refuge (R model). OTUs with a mean intra-OTU divergence lower than 1% were classified as belonging to the R model, and those with higher divergence as belonging to the S model.

### Niche overlap among OTUs and between ecoregions

We quantified niche overlap between informative OTUs with the script script_5_c (supplementary data). Abiotic variables were first reduced to a three-axis PCA space (PC1–PC3). For every informative OTU, we generated a Gaussian hypervolume that describes its climatic niche^58^. Furthermore, we calculated the following pairwise overlap metrics for all possible OTU combinations: Sørensen, Jaccard, and unique-fraction indices. Then, using the random points drawn from each hypervolume as climatic background, following^44^, we built 100 × 100 ecological grids for every OTU pair with the ecospat R package^59^. On each grid we ran 100 permutations of both the niche-equivalence and niche-similarity tests to assess whether observed overlaps differ significantly from expectations under a random-use model^60^.

To evaluate overlap at the ecoregion level we used script_5_d (supplementary data). Each sampling site was assigned to one of three categories: 1) Alone, a single OTU present; 2) Shared, same biome, multiple OTUs present from the same ecoregion; and 3) Shared, different biomes, multiple OTUs present, from different ecoregions. For every informative OTU, we calculated the proportion of its occurrences falling into each category.

### COI molecular clock and Extended Bayesian skyline Plot

To place our analyses in a temporal framework, we derived a molecular clock rate from a well-documented diversification event, the inception of the *Hyalosphenia papilio* species complex. This clade is a group of Arcellinida strictly confined to *Sphagnum* bogs that appeared at the same time as these ecosystems, approximately between 7 and 20 Ma ago^22,23^, in the Miocene. The COI sequences of several lineages (= “cryptic species”) of the *H. papilio* shadow species^12^ were downloaded and analysed in BEAST v1.10.5^61^ to calibrate a molecular clock for Arcellinida. Four independent runs were conducted, each with a different clock model: (1) classical random-local, (2) shrinkage-local, (3) strict, and (4) uncorrelated relaxed. All runs employed a GTR substitution model with estimated base frequencies, and the root age was constrained to 7 Ma using a log-normal prior (mean = 7, SD = 0.1), following^12^. Posterior distributions of *clock.rate*, *branchRatesShrinkage.rate*, and *meanRate* were sampled, discarding the first 20 % as burn-in, and their means were averaged to obtain a single COI substitution-rate estimate for Arcellinida. The resulting rate was compared with published COI clocks for other metazoan phyla (supplementary data S5) and subsequently used to calibrate Extended Bayesian Skyline Plot analyses in BEAST 2 v2.7.7^62^, applying the strict-clock model separately to each informative OTU.

## Acknowledgements

we express our gratitude to A. Berlinches, M. Blázquez, A. García-Bodelón, N. El Khouri-Vidarte, M. Martínez-Ríos, I. Masa-Iranzo, C. Soler-Zamora, Y. Turégano, F. Useros, M. Villar-DePablo, and the MCG lab group for helpful discussions. This work was funded by the Spanish Ministry of Science, Innovation and Universities (MCIN) and the State Research Agency (AEI) under project 20223CNS125, within the 2022 National Plan for Scientific and Technical Research and Innovation (PN2022 - Research Consolidation - State Subprogram for Incorporation - State Programme to Develop, Attract and Retain Talent - PEICTI 2021–2023) / MCIN / AEI / 10.13039/501100011033.S125, as well as under project PID2021-128499NB-I00 State Subprogram for the Generation of Knowledge - State Programme to Promote Scientific and Technical Research. We also acknowledge support to MICIU/AEI /10.13039/501100011033 - FSE+ (JDC2023-050439-I) (RGM).

## Author contributions

R.G.M. contributed Conceptualization, Methodology, Investigation, Visualization, Funding acquisition, Writing - original draft

E. C. contributed in Methodology and revision of the writing

E. L. contributed in Conceptualization, Funding acquisition, Writing - original draft

## Data availability

All data, code, results, including ArKOI metabarcoding database, are available via Figshare (10.6084/m9.figshare.29885813) and github (https://github.com/rubenmiguens/Arcellinida_GLQ). All other data are available in the manuscript or the supplementary materials

